# Expanding the Huntington’s disease research toolbox; validated subdomain protein constructs for biochemical and structural investigation of huntingtin

**DOI:** 10.1101/2022.11.21.516512

**Authors:** Matthew G. Alteen, Justin C. Deme, Claudia P. Alvarez, Peter Loppnau, Ashley Hutchinson, Alma Seitova, Renu Chandrasekaran, Eduardo Silva Ramos, Christopher Secker, Mona Alqazzaz, Erich E. Wanker, Susan M. Lea, Cheryl H Arrowsmith, Rachel J. Harding

**Author notes:** Corresponding Authors, To whom correspondence should be addressed: Rachel J. Harding, Cheryl H. Arrowsmith. Author has updated affiliation: SCIEX, 71 Four Valley Dr, Vaughan, Ontario, L4K 4V8, Canada (Claudia P. Alvarez).

## Abstract

Huntington’s disease is characterised by CAG expansion in the huntingtin gene above a critical threshold of ^~^35 repeats, resulting in polyglutamine expansion of the huntingtin protein (HTT). The biological role of wildtype HTT and the associated mechanisms of disease pathology caused by expanded HTT remain incompletely understood, in part, due to challenges characterising interactions between HTT and putative binding partners. Here we describe a biochemical toolkit of rationally designed, high-quality recombinant HTT subdomains; one spanning the N-terminal HEAT and bridge domains (NTD) and the second spanning the C-terminal HEAT domain (CTD). Using biophysical methods and cryo-electron microscopy, we show these smaller subdomains are natively folded and can associate to reconstitute a functional full-length HTT structure capable of forming a near native-like complex with 40 kDa HTT-associated protein (HAP40). We report biotin-tagged variants of these subdomains, as well as full-length HTT, that permit immobilisation of each protein for quantitative biophysical assays without impacting protein quality. We demonstrate the CTD alone can form a stable complex when co-expressed with HAP40, which can be structurally resolved. The CTD-HAP40 complex binds the NTD, with a dissociation constant of approximately 10 nM as measured by bio-layer interferometry. We validate the interaction between the CTD and HAP40 using a luciferase two-hybrid assay and use subdomain constructs to demonstrate their respective stabilization of HAP40 in cells. These open-source biochemical tools will enable the wider HD community to study fundamental HTT biology, discover new macromolecular or small-molecule binding partners and map interaction sites across this very large protein.

## Introduction

Huntington’s Disease (HD) is an autosomal dominant neurodegenerative disease caused by mutation of the Huntingtin (HTT) gene, resulting in a CAG repeat expansion above a critical threshold of approximately ^~^35 repeats (1). HD patients suffer debilitating cognitive, motor and psychiatric symptoms and have a life expectancy of ^~^18 years after symptom onset (McColgan and Tabrizi, 2018; Saudou and Humbert, 2016). CAG expansion results in an expanded polyglutamine (polyQ) tract near the N-terminus of the encoded protein huntingtin (HTT), resulting in an aberrantly functioning form of the protein (Jimenez-Sanchez et al., 2017; McColgan and Tabrizi, 2018). HTT has been hypothesised to act as a scaffolding protein, governing numerous cellular functions including axonal transport (Gunawardena et al., 2003; Vitet et al., 2020), transcriptional regulation (Ng et al., 2013) and proteostasis regulation by mediating protein-protein interactions (Greco et al., 2022; Rui et al., 2015). Despite identification of the *HTT* gene almost thirty years ago (“A novel gene containing a trinucleotide repeat that is expanded and unstable on Huntington’s disease chromosomes. The Huntington’s Disease Collaborative Research Group,” 1993), there are currently no disease-modifying therapies available, in part due to limited understanding of HTT function, or how these functions might be modulated by polyglutamine expansion. There is therefore a pressing need for the development of chemical and biochemical tools that can be used to illuminate the molecular details of the biological functions of HTT and enable therapeutic strategies to combat the pathology of expanded HTT.

The structure of HTT has been solved by cryo-electron microscopy (Guo et al., 2018; Harding et al., 2021; Huang et al., 2021a), in complex with 40 kDa huntingtin-associated protein (HAP40) (Peters and Ross, 2001), forming a ^~^389 kDa multidomain complex. HTT is primarily composed of HEAT repeats (<u>h</u>untingtin, <u>e</u>longation factor 3, protein phosphatase 2<u>a</u>, and yeast kinase <u>T</u>OR1) (Andrade et al., 2001) that form a large solenoid-like structure at the N-terminal region of the protein as well as a more compact C-terminal region. The N-HEAT and C-HEAT domains are connected by a bridge domain, also composed of HEAT repeats, with HAP40 sandwiched between these regions (**Figure 1A**). HTT is reported to interact with over 3000 proteins (Culver et al., 2012; Greco et al., 2022; Ratovitski et al., 2012; Wanker et al., 2019). The similarity of the HTT-HEAT motifs to related armadillo repeats, which are known to mediate PPIs, may provide a structural basis for HTT interaction network. Additionally, HTT contains many large, disordered regions, including most of exon 1 which contains the polyQ region. Importantly, these disordered regions have been shown to contain post-translational modifications (PTMs) and may mediate interactions with protein binding partners (Ratovitski et al., 2017). HAP40 remains the only biochemically and structurally validated interaction partner (Guo et al., 2018; Harding et al., 2021). Interestingly, while HTT has been recombinantly expressed as an apo protein, efforts to recombinantly express HAP40 on its own have not been successful, suggesting that HTT is required for expression and stabilization of HAP40 structure (Guo et al., 2018; Harding et al., 2021, 2019). This observation has been supported by genetic knockdowns of HTT that show a corresponding decrease in HAP40 expression (Harding et al., 2021; Huang et al., 2021b).

**Figure 1:**
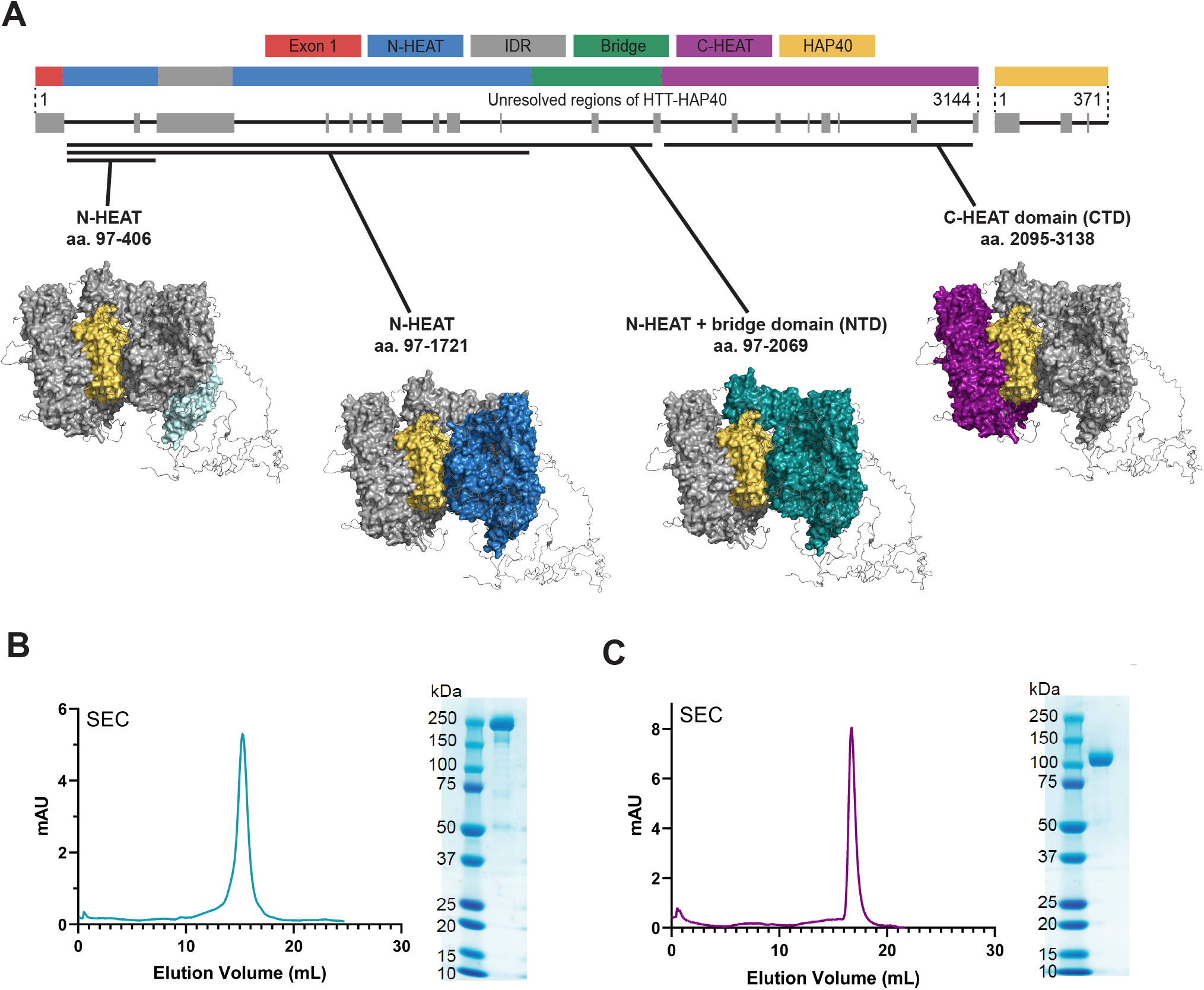
NTD and CTD subdomain constructs of HTT can be recombinantly expressed and produced to high purity in milligram quantities, enabling structural, functional and biophysical studies. A, Schematic representation of the domain architecture of HTT and regions within the domain boundaries that are unresolved in cryo-EM models (grey). Constructs analysed within this study are shown on the HTT-HAP40 structural model (Harding et al., 2021). B, Representative SDS-PAGE and size-exclusion chromatogram of purified NTD. C, Representative SDS-PAGE and size-exclusion chromatogram of purified CTD.

We previously reported a toolkit of resources for the scalable production of high-purity HTT and HTT-HAP40 protein samples from various eukaryotic expression systems (Harding et al., 2019). Milligram quantities of samples can be generated with various Q-lengths as well as an exon 1 deleted (Δ exon 1) form of the protein that are suitable for numerous *in vitro* assays. This expression platform has enabled high-resolution structural analysis of the HTT-HAP40 complex using cryo-EM, and associated biophysical studies such as small-angle X-ray scattering, allowing detailed examination of the structural organisation and potential functional roles of this large multidomain complex (Harding et al., 2021).

Nevertheless, the large size of the protein and its associated HAP40 complex presents challenges for many downstream applications. For example, the analysis of protein-ligand interactions using surface plasmon resonance (SPR), which relies on optical methods to detect mass changes on an immobilised surface, suffer from decreasing signal-to-noise with increasing mass ratio between ligand and analyte molecules (Mitchell, 2010). Smaller subdomain fragments of HTT could serve as useful biochemical tools for the study of HTT function and its interaction with other proteins, permitting a wider array of *in vitro* biochemical and biophysical assays. Additionally, well-defined subdomain fragments could help clarify the relative contributions of each domain in stabilizing HTT-HAP40 interaction, a potential therapeutic target for HD. HTT fragment constructs have been reported in the literature prior to the determination of the cryoEM structure, such as a commonly used fragment encompassing the exon1 region and a portion of the N-HEAT domain (HTT aa. 1-586) (Atwal et al., 2011; Nath et al., 2015). However, these constructs frequently have start and stop sites in the middle of HEAT domains or have boundaries in the middle of extensive disordered regions, which likely result in a non-native protein product. To date, no robust biophysical or structural validation of any such construct has been published.

In this study, we report the cloning, expression, purification and validation of highly pure recombinant HTT subdomain proteins and the open-source tools required to generate these samples. We also report the generation of full-length HTT and HTT subdomain constructs with biotinylated tags at both the N-terminus and C-terminus. We show that constructs encompassing the N-HEAT and bridge domains (NTD, aa. 97-2069) as well as the C-HEAT domain (CTD, aa. 2095-3138), can be stably expressed in eukaryotic cell culture and display good biophysical properties. We demonstrate that these subdomains can be combined *in vitro* or co-expressed to reconstitute the HTT-HAP40 complex, as shown by our cryoEM analysis, indicating that they are folded properly and can serve as suitable surrogates for structural or biophysical analyses. Additionally, we show that HAP40 can form a stable complex with the C-HEAT domain in the absence of the N-HEAT and bridge domains, and that this CTD-HAP40 complex can be resolved by cryoEM. Using these subdomain constructs, we provide the first kinetic analysis of the HTT-HAP40 interaction, revealing high affinity binding of the two proteins. Finally, we validate the interaction between the CTD and HAP40 in cells using a luciferase two-hybrid assay and use subdomain constructs to further probe the HAP40-stabilisation function of HTT in cells. We expect these subdomains will serve as useful biochemical reagents for further structural investigation of the HTT-HAP40 complex, as well as for screening, characterisation, and validation of other members of the HTT protein-protein interactome.

## Results

### Design and cloning of HTT and HTT subdomain constructs with FLAG and Avi tags

Cloning of biotinylated full-length HTT constructs, as well as subdomain constructs, with or without biotin tags, was performed as previously reported (Harding et al., 2019). Ligase-independent cloning (LIC) was used to clone gene sequences into a pBMDEL baculovirus transfer vector which can be used for expression in either Sf9 insect cells or mammalian cells. Clones for subdomain constructs were designed by examining the domain boundaries observed in the cryo-EM model of full-length HTT-HAP40 (**Figure 1A**) and span aa. 97-2069 for the N-HEAT and bridge domain containing construct, and aa. 2095-3138 for the C-HEAT domain construct. All clones were designed to incorporate C-terminal FLAG tags for affinity purification. Biotinylated constructs were cloned to include a 15 amino acid AviTag peptide sequence at either the N- or C-terminus of the protein, enabling biotinylation by co-expression with BirA biotin ligase and addition of exogenous biotin (Fairhead and Howarth, 2015).

### Scalable production of high-purity HTT and HTT subdomain samples

After cloning and verifying the gene sequences of the proposed subdomains, as well as the biotinylated full-length and subdomain constructs, we next aimed to express and purify these constructs using baculovirus-mediated expression in Sf9 insect cell culture. Samples of HTT obtained from either mammalian or insect cell expression have resulted in comparable cryoEM structures (Guo et al., 2018; Harding et al., 2021) and our constructs may be used for production in either system. SDS-PAGE analysis of anti-FLAG purified samples from small-scale test cultures showed the presence of bands corresponding to the expected molecular weights of each subdomain. To further characterise each sample, expression volumes were scaled up to 4 L and the recombinant subdomains were subsequently isolated from lysed cells via FLAG affinity purification and then further purified by size-exclusion chromatography (SEC). We found that a subdomain construct consisting of the folded N-terminal region up to the intrinsically disordered domain (IDR, aa. 97-406) was capable of being expressed and purified by SEC but was found to be poorly stable in biophysical assays (**Figures S1**). Similarly, a construct consisting of the complete N-terminal solenoid region (aa. 97-1721) showed a non-optimal elution profile on SEC, suggesting that it was not a suitable protein for *in vitro* assays (**Figure S2**). However, we discovered that the inclusion of the bridge domain in this construct (aa. 97-2069) caused it to elute as a monodisperse sample, permitting its isolation at >85% purity as determined by densitometry (**Figure 1B**). The yield of the N-terminal HEAT and bridge domain (NTD) construct after this two-step purification sequence was comparable to that of full-length HTT Δexon 1 at ^~^1.4 mg/L. Additionally, a third construct consisting of the C-terminal HEAT domain (CTD) could also be purified as a monodisperse sample in high purity (**Figure 1C**), albeit at lower yields of ^~^0.3 mg/L. Biotin tagged variants of HTT, HTT-HAP40 and the NTD and CTD constructs bearing AviTag sequences at either the N- or C-terminus were co-expressed with BirA and purified in the same manner as constructs bearing only FLAG tags. Size-exclusion chromatography profiles and corresponding yields of biotinylated proteins were nearly identical to non-biotinylated constructs. The installation of the biotin moiety was confirmed by streptavidin gel-shift (Fairhead and Howarth, 2015) (**Figure S3**).

### Subdomain constructs show good biophysical properties in multiple orthogonal assays

With purified samples of each subdomain in hand, we next sought to assess their quality and characteristics through a series of biophysical analyses. We performed differential scanning fluorimetry (DSF) (Niesen et al., 2007) to determine the thermal stability of each construct under identical buffer, salt and pH conditions as full-length HTT samples (**Figure 2A-2C**). We found that the NTD, with a calculated melting temperature (T_m_) of 48.3 ± 0.2°C, has significantly greater thermal stability relative to full-length apo HTT (T_m_ ^~^43.2 ± 0.1°C). Conversely, we observed a T_m_ of 42.4 ± 0.1°C for the CTD, suggesting this subdomain is less thermostable than full-length HTT. Nevertheless, both subdomains showed desirable biophysical characteristics, including a sharp transition upon heating and high dynamic range of the fluorescence signal between folded and unfolded states (**Figure S4**). We also analysed the protein constructs using differential static light scattering (DSLS) to assess the aggregation behaviour of each sample upon heating (**Figure 2D-2F**). Mirroring the trend observed by DSF, the NTD showed a significantly higher aggregation temperature (T_agg_) than the CTD. Given the large, multidomain structure of full-length HTT, it is possible that its T_m_ and T_agg_ values represent an averaging of the values for the NTD and CTD. Notably, the unfolding of full-length HTT as monitored by DSF occurs over a broader temperature range, suggesting that the domains may unfold sequentially. Taken together, these data suggest that the two purified subdomains each comprise a soluble, folded protein that is suitable for use in biophysical and screening assays.

**Figure 2:**
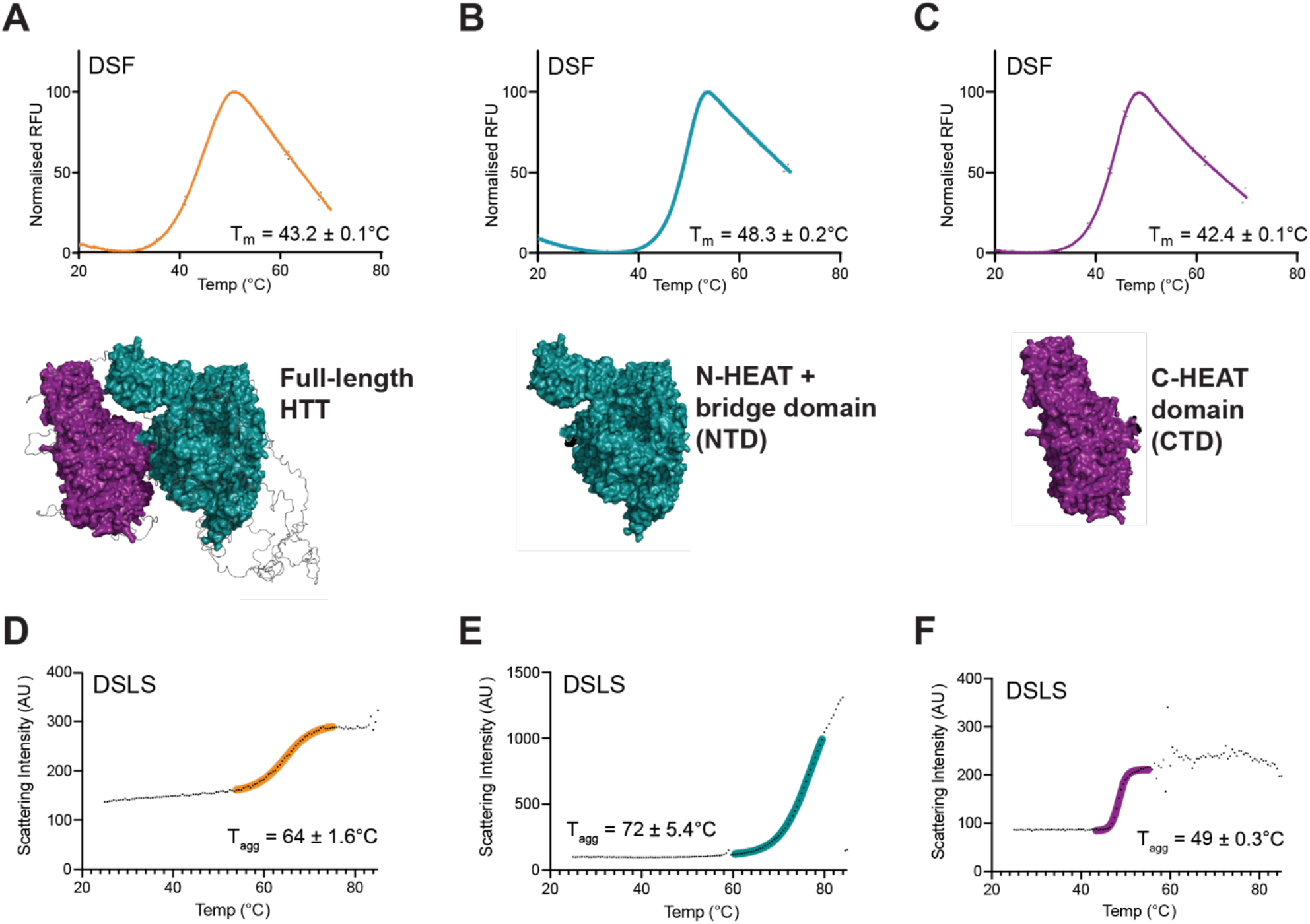
NTD and CTD subdomains of HTT have good biophysical properties. A-C, DSF profiles and calculated melting temperature (T_m_) values for full-length HTT (orange) vs NTD (teal) and CTD (magenta). Melting temperatures were determined from the inflection point of curves obtained by fitting the data to the Boltzmann sigmoidal function. D-F, DSLS profiles of full-length HTT (orange) *vs*. NTD (teal) and CTD (magenta) showing calculated aggregation temperature (T_agg_).

### CryoEM reveals the HTT subdomains can stabilise HAP40, forming a functional HTT-HAP40 complex

Given the results of our biophysical assessment of the NTD and CTD samples, which suggested the proteins were properly folded, we next examined if they could be used to re-constitute the HTT-HAP40 complex. To test this, we co-expressed NTD, CTD, and HAP40 in Sf9 cells and attempted to purify the protein as before. Strikingly, after affinity purification from lysed cells with anti-FLAG resin, we observed protein bands corresponding with the molecular weight of these three fragments by SDS-PAGE (**Figure S5**). Further purification of the crude samples by size-exclusion chromatography as before revealed a profile nearly identical to that obtained with full-length HTT-HAP40, with a major peak eluting at the same volume as expected for the intact construct (**Figure 3A**). After collecting this peak and performing further analysis by DSF, we observed a thermal transition profile and T_m_ value nearly identical to that of full-length HTT (**Figure 3B, Figure S6**). As further confirmation of the protein bands observed through SDS-PAGE analysis of the purified triplet complex (**Figure 3C**), we carried out western blotting using antibodies for the N-terminal region of HTT as well as HAP40 (**Figure 3D, Figure S7**). This analysis showed that the 218 kDa band corresponds to the NTD and that the band at ^~^40 kDa is HAP40. We also noted the presence of an additional band at ^~^-160 kDa, which persisted despite several rounds of purification. This impurity also possessed a recognised epitope of the D7F7 α-HTT antibody, suggesting that it may be a partial fragment of the N-HEAT region that either does not fully express in Sf9 cells or has been proteolysed by an unknown mechanism. HTT is susceptible to proteolysis within the IDR (30), and the molecular weight of this fragment suggests this band corresponds to a cleavage product within this region.

**Figure 3:**
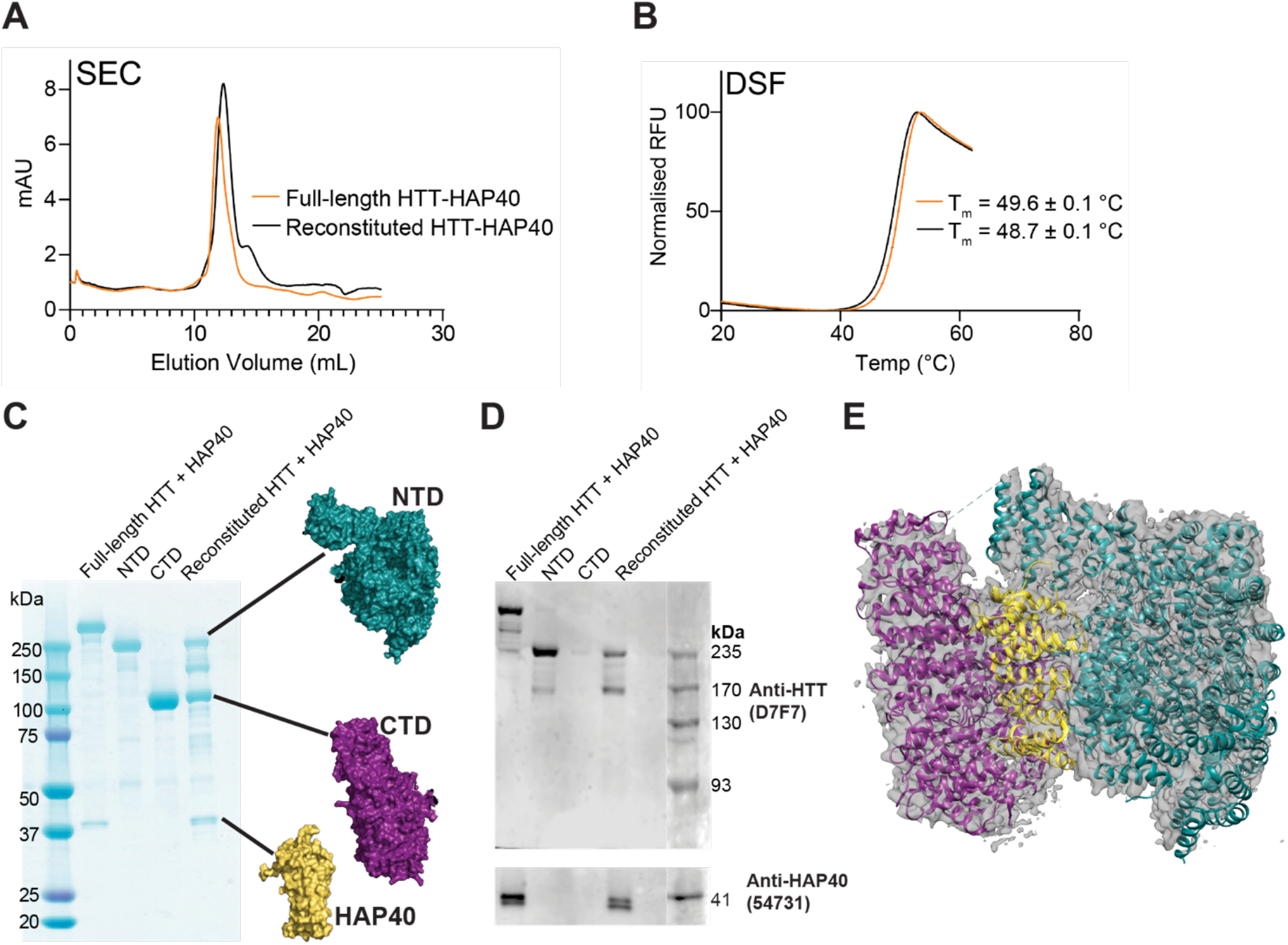
HTT subdomains bind HAP40 and can be used to reconstitute the HTT-HAP40 complex. A, Size exclusion chromatography profiles of co-expressed NTD, CTD and HAP40 (black trace) after FLAG purification from Sf9 cells. An elution volume nearly identical to full-length HTT-HAP40 (orange trace) indicates formation of the trimeric complex. B, Reconstituted and full-length HTT-HAP40 complexes have similar melt profiles as determined by DSF. C, SDS-PAGE of full-length HTT (lane 1), NTD (lane 2), CTD (lane 3) and co-expressed subdomains (lane 4). D, Western blots of full-length HTT-HAP40, NTD, CTD, and reconstituted HTT-HAP40. E, Overlay of cryo-EM map obtained from reconstituted HTT-HAP40 superimposed on the model generated for full-length HTT-HAP40 6X9O.

To further verify the structural fidelity of the reconstituted complex, we performed cryo-EM of this sample using similar conditions as performed for the full-length HTT-HAP40 complex (**Table S1**). The 3.3 Å resolution map obtained from this sample is highly similar to the previously solved atomic resolution model (**Figure 3E**) (Harding et al., 2021). The striking resemblance of this reconstituted complex to the full-length HTT-HAP40 complex indicates the subdomains are capable of properly folding and associating to a degree that enables the concomitant expression of HAP40 and validates these constructs for the expression of functional and folded HTT protein subdomains.

### The HTT CTD is sufficient to stabilise HAP40, forming a complex which can be resolved by cryoEM

Having confirmed the robust structural and biophysical integrity of the NTD and CTD constructs, we next asked whether either the NTD or CTD alone is capable of stabilizing HAP40 when co-expressed. Co-expression experiments with the NTD and analysis by SDS-PAGE did not show evidence of HAP40 expression (**Figure S8**). However, after co-expressing HAP40 with the CTD, a band corresponding to HAP40 was present on SDS-PAGE at the expected molecular weight (**Figure 4A, Figure S9**). Analysis of this sample using analytical size-exclusion chromatography revealed a significant shift in elution volume, corresponding to an increase in size relative to the C-HEAT construct (**Figure 4B**). We performed DSF to determine the thermostability of this complex and found that it possessed a marginally higher melting temperature of approximately 1.6 degrees and a sharper melting transition curve than the CTD (**Figure 4C, Figure S10**). This suggests that the presence of the HAP40 binding partner is beneficial to the overall stability of the CTD and that the complex is properly folded.

**Figure 4:**
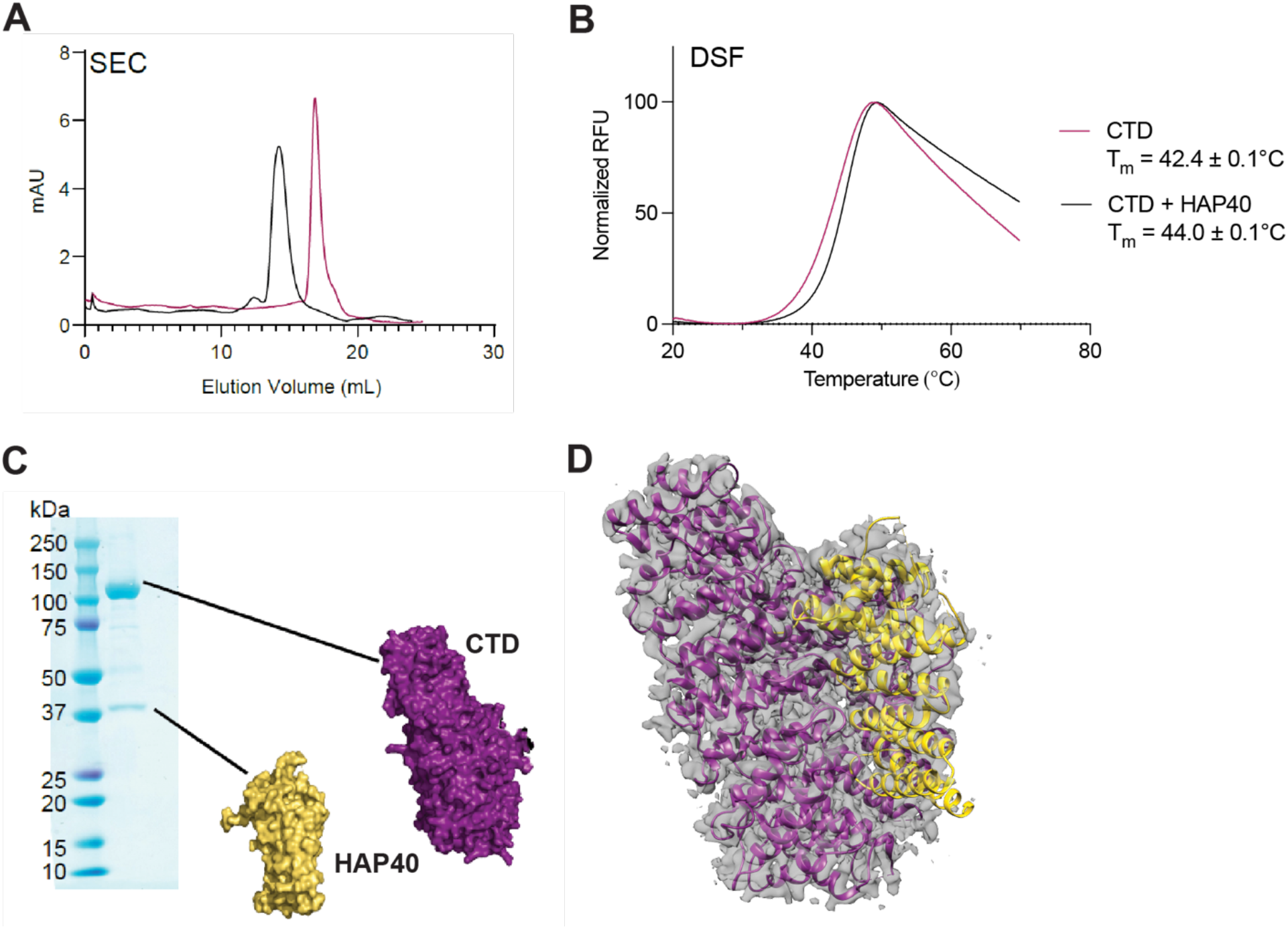
HAP40 can be expressed with CTD alone. A, Overlay of size-exclusion chromatograms of purified co-expressed CTD and HAP40 (black) vs apo CTD (magenta). B, SDS-PAGE analysis of purified co-expressed C-HEAT and HAP40, with bands at ^~^120 kDa and ^~^40 kDa indicated. C, The CTD-HAP40 complex displays higher thermostability and a sharper transition upon thermal unfolding as determined by DSF. D, Overlay of the map of purified co-expressed CTD-HAP40 obtained by cryo-EM (grey mesh) with the carton structure of the domains determined from the full-length cryo-EM model 6X9O.

We again studied the structure of this complex by cryo-EM to determine its overall 3D structure relative to full-length HTT-HAP40 (**Figure 4D, Table S1**). We found that the 3.2 Å resolution map obtained from the CTD-HAP40 complex fit well into the structural model derived from the full-length complex for these regions of the protein, with only slight deviations at the termini of the proteins. Overall, these data indicate that the CTD alone is sufficient to stabilize and permit expression of HAP40.

### HTT subdomain constructs allow interrogation of protein-protein interactions by biophysical methods

With the CTD-HAP40 complex in hand, we proceeded to measure its binding affinity to the NTD *in vitro*. Using analytical size-exclusion chromatography, we showed that pre-mixing the purified protein subdomains results in a significant leftward shift in elution volume compared to the elution of CTD-HAP40 alone, suggesting that the majority of the CTD-HAP40 associates with the NTD (**Figure 5A**). To quantify the interaction, we purified CTD-HAP40 bearing a C-terminal biotin tag to measure binding kinetics using biolayer interferometry (BLI). Biotinylated CTD-HAP40 was immobilised onto streptavidin biosensors at a concentration of 1 μg/mL and, after equilibration, the sensors were incubated with various concentrations of NTD (**Figure 5B, Figure S11**). The resulting association and dissociation curves revealed a concentration-dependent rate of binding and dissociation of the two protein subunits. By fitting the curves to a global 1:1 binding model, a dissociation constant (Kd) of 10 ± 0.3 nM was obtained. These data represent the first quantitative *in vitro* assessment of the strength of HAP40 binding to HTT and support earlier observations that the HTT-HAP40 interaction is remarkably stable under a wide array of conditions (Harding et al., 2021).

**Figure 5:**
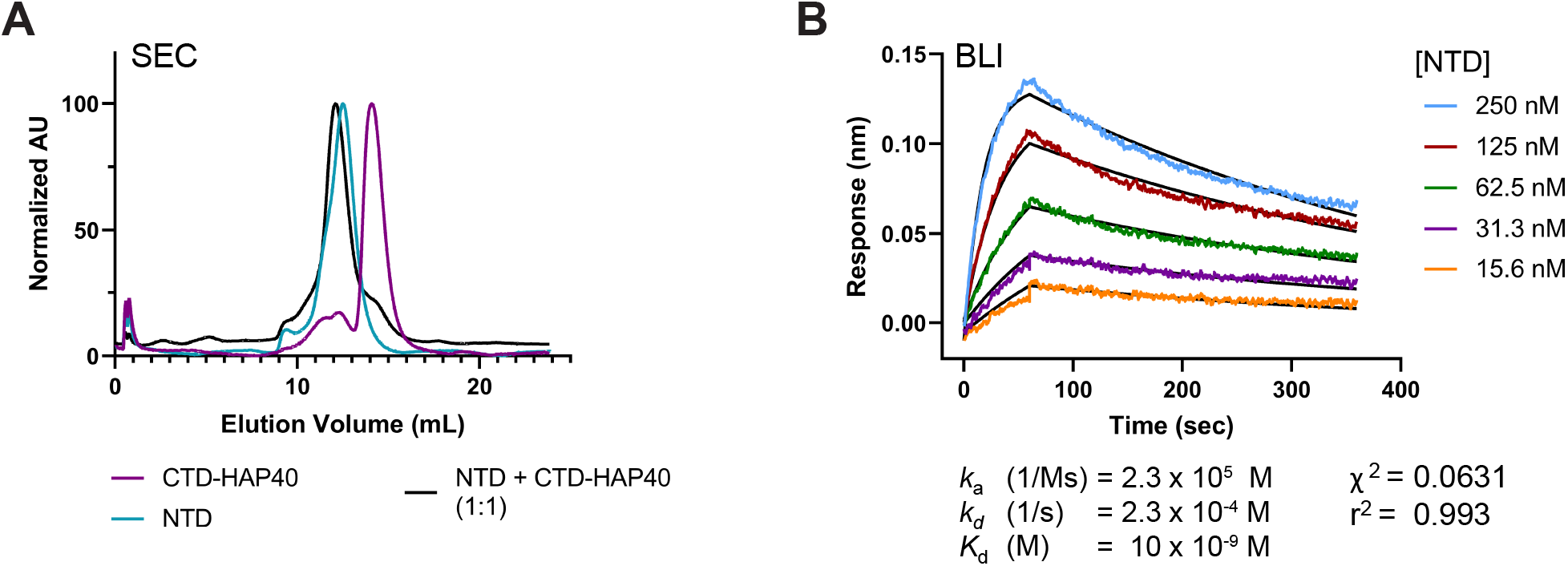
Purified co-expressed CTD-HAP40 binds the NTD with high affinity. A, Overlayed size-exclusion chromatography profiles of purified CTD-HAP40 (magenta), NTD (teal) and equimolar amounts of pre-mixed NTD and CTD-HAP40 (black). B, Characterisation of NTD binding affinity to immobilised CTD-HAP using biolayer interferometry, revealing a Kd of 10 ± 0.3 nM. Kinetic parameters were determined by fitting the data to a 1:1 binding association and dissociation model (black lines) using GraphPad Prism 9. Three independent experiments were performed.

### Validation of HTT subdomain protein-protein interactions in cells

To further verify that HAP40 can be bound by HTT CTD to form a stable complex under physiological conditions, we performed a luciferase two-hybrid (LuTHy) assay (Trepte et al., 2018) in HEK293 cells (**Figure 6A**). mCitrine-Protein A (mCit-PA)-conjugated HTT constructs corresponding to the full-length and CTD sequences, showed BRET ratios which were significantly increased compared to tag-only controls when co-expressed with HAP40-NanoLuc luciferase (NL) fusion protein (**Figure 6B**). This confirms that the CTD of HTT is sufficient to bind HAP40 in a cellular context.

**Figure 6:**
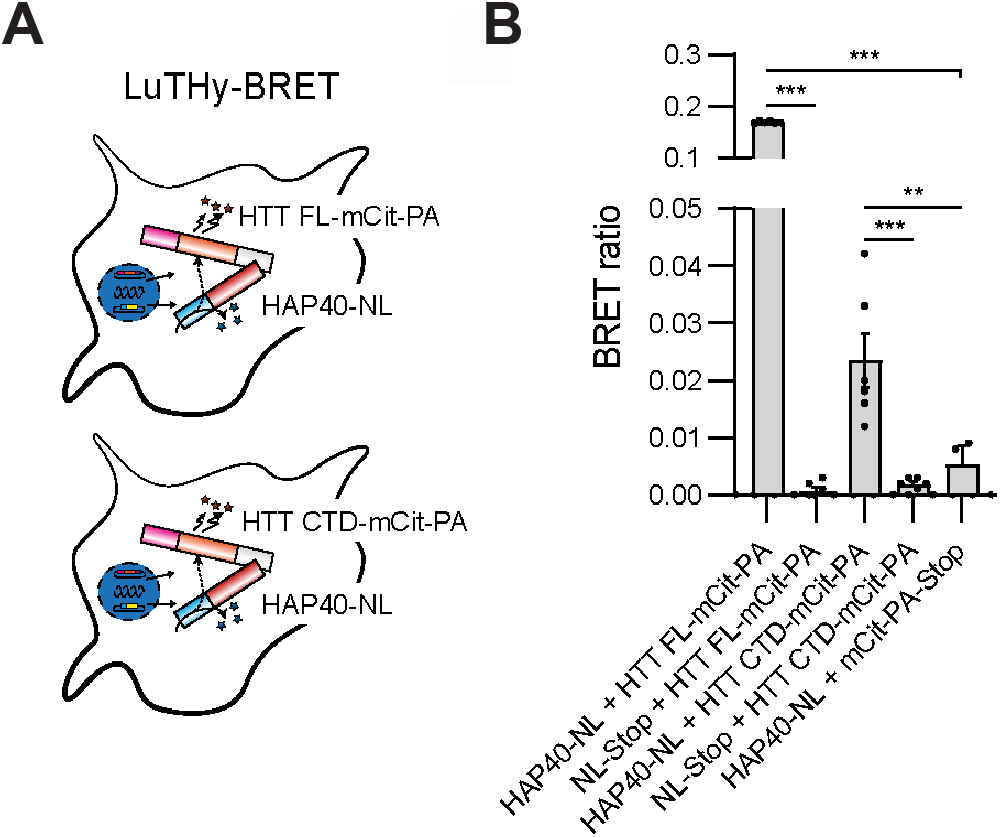
LuTHy assay shows interaction of HTT full-length (FL) and its C-terminal domain (CTD) with HAP40 in live cells. A, Graphical illustration of the LuTHy-BRET assay performed. HTT FL or HTT CTD were expressed as mCitrine-Protein A (mCit-PA)- and HAP40 as NanoLuc luciferase (NL)-tagged fusion proteins in HEK293 cells. After expression for 48 h and addition of luciferase substrate, BRET was quantified from live cells. B, BRET ratios between HAP40-NL and HTT FL-mCit-PA and HTT CTD-mCit-PA. As a control, NL only was co-transfected with the HTT acceptor constructs, respectively, as well as PA-mCit only with the HAP40-NL donor construct. HAP40-NL co-expressed with HTT FL-mCit-PA and HTT CTD-mCit-PA showed significantly increased BRET ratios compared to controls, respectively. Bars represent means ± SEM from two independent experiments performed in triplicate. One-way ANOVA with Tukey’s multiple comparisons test, **p < 0.002, *** p < 0.001.

### Investigating HTT cellular functions with subdomain constructs

Next, we sought to scrutinise a defined biomolecular function of HTT using our subdomain tools. HAP40 expression cannot be detected in HTT knockout HEK293 cells (**Figure 7A**), in line with previously published data which show the levels of HAP40 are dependent on HTT protein levels and that HTT functions to stabilise HAP40 (Harding et al., 2021; Huang et al., 2021b). Thus, by monitoring the protein levels of HAP40 in HTT KO cells in the presence of exogenous full-length or subdomain constructs of HTT we could observe a rescue of the HAP40 null phenotype in these cells. Overexpression of N-HEAT, CTD or full-length HTT in these cells showed that only the full-length HTT can restore some degree of HAP40 expression in HTT knock out HEK293 cells. Together with our LuTHy assay data, this indicates that although HTT CTD can bind HAP40 in a cellular context, it is not sufficient to restore the HAP40 stabilisation function of HTT.

**Figure 7:**
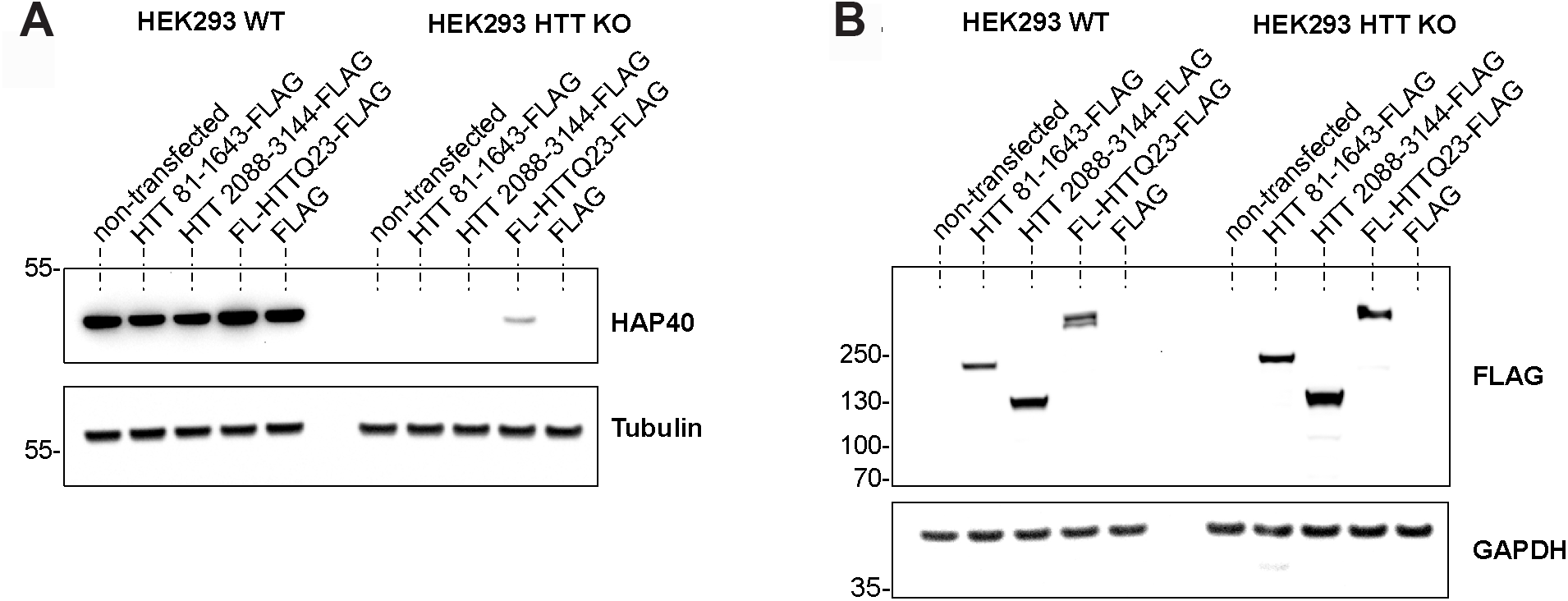
Only overexpression of full-length HTT can rescue HAP40 protein levels in HTT-null cells. A-B, Representative western blot images of total cell lysates from HEK293 wildtype and HEK293 HTT-null lines after transfection with of various FLAG-tagged constructs for HTT full-length (FL) and truncated species of HTT. A, Nitrocellulose membrane was incubated with rabbit anti-F8A1 antibody, followed by mouse anti-tubulin to detect endogenous HAP40 and tubulin. B, Nitrocellulose membrane was incubated with mouse anti-FLAG antibody, followed by mouse anti-GAPDH to detect overexpressed FLAG-tagged constructs and endogenous GAPDH. Calculated molecular weight for HTT 81-1643 is 175 kDa, for HTT 2088-3144 121 kDa, and for FL HTTQ23 352 kDa. Two independent experiments were performed.

## Discussion

The expansion of the HTT polyQ tract has been well-defined as the cause of HD pathobiology. However, the precise biochemical functions of wildtype HTT, or the changes that result in the presence of expanded HTT, remain incompletely understood. Given a lack of biochemical tools to interrogate HTT function in cells, genetic studies have formed the basis of most functional studies, either through knockdown of HTT expression or the generation of expanded polyQ allelic series in various animal models or cell lines (Menalled et al., 2009; Shen et al., 2019; Southwell et al., 2013). Additionally, affinity-based proteomic methods coupled with mass spectrometry have been applied to identify putative protein binders that interact with HTT (Culver et al., 2012; Ratovitski et al., 2012). Efforts to interrogate HTT function by many biochemical and biophysical assays are thwarted by the large size of HTT, which renders these assays non-tractable. Smaller HTT subdomains provide a solution to this problem, but many of the constructs published to date are not designed with knowledge of the HTT structure in mind, meaning their boundaries may interrupt structural elements or they contain extensive disordered regions, which likely result in non-native protein products. To date, no robust biophysical or structural validation of any such construct has been published to date. The toolkit of subdomain constructs described in this manuscript provides a solution to this problem, through the rational structure-based design of subdomain constructs and their validation by numerous assays and structural analysis by cryoEM.

Because of the suspected role of expanded HTT in triggering HD progression, HTT lowering therapies have been hypothesised as a potential therapeutic strategy for HD. One proposed pharmacological approach to HTT lowering is through the use of proteolysis-targeting chimeras (PROTACs) – heterobifunctional ligands that combine a recognition scaffold for a protein of interest combined with an E3 ligase motif, resulting in ubiquitin-mediated proteasomal degradation of target proteins (Sakamoto et al., 2001; Schapira et al., 2019). To date, however, all small-molecule ligands being developed as PROTACs bind mutant aggregated HTT fragments (Hirai et al., 2022; Tomoshige et al., 2017). As a result, these compounds may suffer from off-target effects because they do not bind a homogenous or specific binding site on the protein and do not bind soluble HTT that is present within cells. There is therefore a need for the discovery and validation of novel small-molecule scaffolds that bind soluble HTT in a reversible and specific manner.

To support efforts to validate the complex network of HTT protein-protein interactions, as well as to support pre-clinical characterisation and optimisation of small-molecule ligands, high quality biochemical tools and assay methods are needed. The suite of full-length and subdomain constructs we have developed have a range of potential applications to support fundamental biology and drug discovery efforts. We expect that the combined use of NTD and CTD constructs will permit approaches to counter-screen novel ligands for selectivity of binding, as well as to characterise the degree and site of interaction between HTT and potential PPIs. Furthermore, the smaller size and robust nature of the biotin tags on these constructs enable a variety of biophysical assays to accurately quantify the affinity of potential HTT binders. Additionally, the smaller molecular weight of the subdomains and, in some cases, their higher expression yield relative to full-length HTT, improves their suitability for high-throughput applications.

Our efforts to generate truncated HTT subdomains have also provided fundamental insight into the key structural interactions governing the HTT-HAP40 complex. Our observation that HAP40 can be co-expressed with the CTD alone, both in vitro, and form a stable, soluble protein complex, shows the importance of this interface for HAP40 stability. Analysis of the CTD-HAP40 interaction surface shows that this subdomain accounts for approximately 60% of the total interaction area between HAP40 and full-length HTT, highlighting the significance of the C-HEAT on binding to HAP40. The generation of this complex permitted quantitative characterisation of its binding to the NTD, providing additional evidence of the robust nature of HTT-HAP40 affinity. Such analysis has not been possible with full-length HTT to date as HAP40 must be coexpressed with HTT, or, as we demonstrate in this study, the CTD of HTT. Given that no other putative protein binding partners of HTT have been characterized through a direct binding assay in this manner, the use of the CTD-HAP40 and NTD association provides an important positive control for prospective PPI analyses.

Using our defined and validated subdomain construct boundaries, we were also able to characterise the CTD-HAP40 interaction in live cells. This revealed that the CTD can bind HAP40 when measured using the BRET-based LuTHy readout. Interestingly, in a HTT knockout background where HAP40 protein levels are ablated due to lack of stabilisation by HTT (Harding et al., 2021; Huang et al., 2021b; Xu et al., 2022), we showed that the full-length, but not the CTD alone, is able to restore HAP40 expression. This suggests that the interdependent relationship between HTT and HAP40 expression is perhaps not just one of structural stabilisation, as the CTD is sufficient to form a stable HAP40 complex, but perhaps linked in a more complex co-translational dependent manner.

In summary, we have generated and validated a suite of high-quality biochemical tools to support the structural and functional characterisation of HTT. We report the design and purification of HTT subdomains encompassing the N-terminal and C-terminal regions of the protein, as well as biotinylated variants that permit convenient immobilisation on streptavidin medium. These protein constructs, which are stably expressed and folded, permit a wide array of biophysical assays to probe the diverse network of HTT protein-protein interactions and facilitate future screening efforts. Additionally, these subdomains have provided important information on the nature of the HTT-HAP40 interaction, demonstrating that the C-HEAT domain forms key interactions with HAP40 and is sufficient to enable the expression of HAP40 in the absence of the NTD, both *in vitro* and in cells. Finally, we used our defined subdomains to probe the HAP40-stabilisation function of HTT, revealing expression of the full-length HTT protein molecule is required to rescue this function in a knockout background. We expect that these biochemical tools will find widespread use within the HD community and will support ongoing efforts to elucidate the biological roles of HTT.

## Methods

### Cloning of expression constructs

Expression constructs for the NTD (amino acids 97 to 2069) and the CTD (amino acids 2095 to 3138) were constructed with three different affinity tag arrangements. The first version had a C-terminal Flag tag, the 2nd version had a N-terminal biotinylation tag / C-terminal Flag tag, and the 3rd version had the biotinylation and Flag tag on the C-terminal. The biotinylation tag contains the sequence GLNDIFEAQKIEWHE which facilitates the in vivo conjugation of a single biotin to the lysine residue when the HTT protein is co-expressed with biotin ligase in SF9 cells. To assemble each plasmid, the DNA encoding for the HTT gene fragments was PCR amplified from cDNA (Kazusa clone FHC15881). PCR primers were designed to add coding sequence for N-terminal or C-terminal biotinylation tags and Flag tags as required. The HTT expression constructs were assembled using the BD In-Fusion PCR cloning kit in the mammalian/insect cell vector pBMDEL (Addgene plasmid #111751), an unencumbered vector created for open distribution of these reagents.

Two expression constructs were made for the expression of full length (FL) HTT (amino acids 1 to 3144). One had an N-terminal biotinylation tag and C-terminal flag tag. The other had both the biotinylation tag and Flag tag on the C-terminal. These plasmids were created through modification of Addgene plasmid #111726, whose construction we previously described (Harding et al., 2019). Briefly, these modifications were done in two steps. First, a N-terminal or C-terminal fragment the HTT gene with the required tags was assembled into an intermediate plasmid. Second, these tagged fragments were transferred into Addgene plasmid #111726 by PCR amplification followed by insertion into the expression construct using the BD In-Fusion PCR cloning kit.

Expression vectors for C-terminal FLAG-tagged constructs were generated by Gateway cloning. To this end, 150 ng of entry vector in a pDONR221 vector for full-length HTTQ23, HTT 81-1643, and HTT 2088-3144 was mixed with 600 ng of the destination vector pDEST_gateway-2xFLAG (Addgene plasmid #118372), 1 μl of LR clonase, and filled up to 10 μl with TE buffer pH 8.0. Reaction was incubated overnight at room temperature. Afterwards 1 μl of Proteinase kinase K was added and incubated at 37°C for 10 minutes. Reaction was then transformed into chemically competent Mach1 bacterial cells, plated onto ampicillin agar plates. Clones were picked, cultured in liquid LB medium, and DNA isolated using QiAprep Spin Miniprep Kit (Qiagen). Final constructs were sequenced to validate correct insertion using the following primers; GAGGTCTATATAAGCAGAGC and AACCATTATAAGCTGCAATAAAC.

LuTHy plasmids were generated as described previously (Trepte et al., 2018). In brief, open reading frames (ORFs) of full-length HTT(Q23) or HTT’s CTD were amplified from previously generated expression plasmid (Addgene plasmid #111723) and resulting attB1 and attB2-flanked PCR products were subcloned into pDONR221 entry vectors using BP Clonase (Gateway Cloning System, Invitrogen). For HAP40 (F8A1), a cDNA entry clone was obtained from Source BioSciences (OCAAo5051D1091D). To generate LuTHy expression plasmids, HAP40 was cloned into a LuTHy donor vector (Addgene plasmid #113447) and the HTT constructs into a LuTHy acceptor vector (Addgene plasmid #113449) by LR Clonase reactions (Gateway Cloning System, Invitrogen). All expression plasmids were finally validated by restriction enzyme digest, agarose gel electrophoresis, and Sanger sequencing.

The DNA sequences and expressed protein sequences of plasmids used in this paper are summarized in **Appendix 1**.

### Protein expression and purification

Expression of HTT subdomains was performed in Sf9 insect cells as previously described (Harding et al., 2019) with minor modifications. Briefly, cells were infected with P3 recombinant baculovirus and grown at 37°C until cell viability reached 80-85%, normally ^~^72 hours post-infection. For full-length HTT-HAP40, a ratio of 1:1 HTT:HAP40 P3 baculovirus was used for infection. For CTD-HAP40, a ratio of 1:3 CTD:HAP40 P3 baculovirus was used. Biotinylation of AviTag constructs was achieved by co-expression with recombinant BirA in the presence of μM biotin. For purification, cells were harvested by centrifugation and lysed with two freeze-thaw cycles. Lysates were then clarified by centrifugation at 29, 416 x g for 60 minutes. After collecting the supernatant, proteins were isolated by FLAG affinity chromatography. Crude protein samples from FLAG-eluted fractions were then pooled and purified by size-exclusion chromatography using a Superose 6 Increase 10/300 GL column (Cytiva Life Sciences) in buffer containing 20 mM HEPES pH 7.4, 300 mM NaCl, 2.5% (v/v) glycerol, 1 mM TCEP. Peaks corresponding to monodisperse protein were pooled, concentrated and flash frozen and stored at −80°C until use. Sample purity was assessed by SDS-PAGE.

### Differential scanning fluorimetry

Determination of protein thermostability by DSF was performed using a Roche Applied Science Light Cycler 480 II. Samples were prepared in LightCycler 480 white 384-well plates (Roche) at a volume of 20 μL per well using a final concentration of 0.1 mg/mL protein and 5X Sypro Orange (Invitrogen). All samples were diluted in buffer containing 20 mM HEPES pH 7.4, 300 mM NaCl, 2.5% (v/v) glycerol, and 1 mM TCEP. Thermal shift assays were carried out over a temperature range of 20 to 95°C at a ramp rate of 0.02°C/sec. Fluorescence measurements were taken using a 465 excitation / 580 emission filter set. Reactions were performed in triplicate and data were analyzed using GraphPad Prism 9. Melt curves were fitted to a Boltzmann sigmoidal curve and apparent T_m_ values were determined by calculating the inflection point of the fitted curves.

### Differential static light scattering

T_agg_ values for purified proteins were determined by DSLS using a Stargazer instrument (Harbinger Biotech). Protein samples were diluted to 0.4 mg/mL in buffer containing 20 mM HEPES pH 7.4, 300 mM NaCl, 2.5% (v/v) glycerol and 1 mM TCEP using a volume of 50 μL per well. Samples were heated from 20°C to 85°C at a rate of 1°C/min and protein aggregation was monitored by measuring the intensity of scattered light every 30 s with a CCD camera. Scattering intensity was then plotted and fitted to the Boltzmann equation, and T_agg_ values were determined by measuring the inflection point of each curve.

### Cryo-EM sample preparation and data acquisition

Full-length HTT-HAP40 or CTD-HAP40 samples were diluted to 0.2 mg/mL in 25 mM HEPES pH 7.4, 300 mM NaCl, 0.025% w/v CHAPS and 1 mM DTT and adsorbed onto gently glow-discharged suspended monolayer graphene grids (Graphenea) for 60 s. Grids were then blotted with filter paper for 1 s at 100% humidity, 4 °C and frozen in liquid ethane using a Vitrobot Mark IV (Thermo Fisher Scientific).

Data were collected in super-resolution counting mode on a Talos Arctica (Thermo Fisher Scientific) operating at 200 kV with a BioQuantum imaging filter (Gatan) and K3 direct detection camera (Gatan) at 100,000x magnification, corresponding to a real pixel size of 0.81 Å/pixel. Movies were collected at a dose rate of 22.8 e^-^/Å^2^/s, exposure time of 2.25 s, resulting in a total dose of 51.3 e^-^/Å^2^ fractionated across 53 frames.

### Cryo-EM data processing

Movies were processed in real time using the SIMPLE 3.0 pipeline (Caesar et al., 2020), using SIMPLE-unblur for patched motion correction, SIMPLE-CTFFIND for patched CTF estimation and SIMPLE-picker for particle picking. Particles were extracted in 288 × 288 pixel boxes, sampling of 0.81 Å/pixel.

For full-length HTT-HAP40, 112,717 particles were generated from 166 movies and subjected to 2D classification (50 classes, 160 Å mask) within cryoSPARC (Punjani et al., 2017). Particles (64,597) belonging to the most defined, highest resolution classes were used to generate an ab initio map. This map was lowpass filtered to 30 Å and used as reference for non-uniform refinement within cryoSPARC, yielding a 3.3 Å volume as assessed by Gold standard Fourier Shell Correlations (FSC) using the 0.143 criterion.

For CTD-HAP40, 986,229 particles were generated from 1,461 movies and subjected to initial 2D classification in SIMPLE 3.0 (200 classes, 140 Å mask). “Good” particles (819,568) were subjected to an additional round of 2D classification within cryoSPARC (200 classes, 140 Å mask). Retained particles (297,261) underwent multi-class (4 classes) ab initio volume generation within cryoSPARC, producing one sensible class composed of 134,849 particles. The volume corresponding to this class was lowpass filtered to 30 Å and used as reference for non-uniform refinement within cryoSPARC, generating a 3.2 Å volume as assessed by Gold standard Fourier Shell Correlations (FSC) using the 0.143 criterion.

### Western blotting of recombinant protein samples

General protocols for western blot analyses were performed as previously described (Harding et al., 2021). Primary antibodies used include anti-HTT D7F7 (Cell Signalling Technology), anti-HAP40 54731 (Novus Biologicals) and anti-FLAG F1804 (Sigma). Secondary antibodies used are goat-anti-rabbit IgG-IR800 (LI-COR) and donkey anti-mouse IgG-IR680 (LI-COR). Membranes were visualised using an Odyssey CLx imaging system (LI-COR).

### Biolayer interferometry

The affinity of the CTD-HAP40 binding interaction to the NTD was determined using an OctetRED96 BLI system (ForteBio). Experiments were performed at 25°C in 96-well black microplates (Greiner, 655209) with shaking at 1000 rpm. Purified CTD-HAP40 with a C-terminal biotin tag was diluted to 1 μg/mL in buffer containing 100 mM sodium phosphate pH 7.4, 500 mM NaCl, 0.05% Tween-20, and 0.1% BSA and were loaded onto streptavidin biosensors (ForteBio, 18-5019) for 300s. Sensors were then washed in buffer for 120 s to establish a baseline reading. A serial dilution of NTD in identical buffer was prepared in the same plate, and then loaded sensors were transferred into NTD wells for a 60 s association period to determine k_on_. Sensors were then transferred to wells containing only buffer for 300 s to determine k_off_. Binding kinetics were determined using GraphPad Prism 9 by fitting the data to a 1:1 association then dissociation model. Association and dissociation curves from all concentrations of NTD were fitted globally to a shared k_on_ and k_off_ value. Background signal from reference sensors without immobilised CTD-HAP40 was subtracted from experimental sensor wells.

### Cell culture and transfection

Human embryonic kidney line 293 (HEK293) wildtype and HTT-null cells were grown in Dulbecco’s modified Eagle’s medium (Thermofisher #41965) supplemented with 10% heat-inactivated fetal bovine serum (ThermoFisher #10500064), and 1% penicillin/streptocillin (ThermoFisher #15140122) at 37 °C, and 5% CO2. Cells were subcultured every three to four days. For overexpression studies, one million cells in a final volume of 2 ml complete DMEM medium were reversed transfected in 6-well plates with a transfection mix composed of two μg of DNA in a volume of 200 μl Opti-MEM and 5 μl Fugene transfection reagent (2:5:1 ratio). After 48 hours cells were trypzined, washed with ice-cold PBS, pelleted and stored at −80°C.

### Western blotting of cell lysates

HEK293 cell pellets collected from a 6-well plate were lyzed in 30-50 μl HEPES lysis buffer (50 mM HEPES pH 7.0, 150 mM sodium chloride, 10 % glycerol, 1 % NP-40, 20 mM NaF, 1.5 mM MgCl2, 1 mM EDTA, 1 mM PMSF, 0.5% sodium deoxycholate, 1x Benzonase, 1x, Roche Complete EDTA-free protease inhibitor cocktail (Merck, 5056189001)) for 30 min on ice. Lysates were centrifuged at 14,000 rpm for 10 min at 4°C and supernatants collected. Protein concentrations were determined using the PierceTM BCA assay (Thermo Scientific) and 20 μg total protein was combined with 50 mM DTT and 1x NuPAGE LDS sample buffer, followed by 5 min at 95°C. Proteins were separated by SDS-PAGE using a NuPAGE 4-12% Bis-Tris gel and transferred onto nitrocellulose membranes (Cytiva, 10600002). Membranes were blocked for 1 hour in 3% milk in PBS with 0.05% Tween. The following primary antibodies were applied overnight at 4°C: mouse anti-FLAG (1:1000; Sigma #F3165), rabbit anti-HAP40 (1:1000; Atlas Antibodies #HPA046960), mouse anti-tubulin (1:80,000; Sigma #T6074), and mouse anti-GAPDH (1:1000; Santa Curz #sc-47724). The following secondary antibodies diluted to 1:6000 in 3% milk and applied for two hours at room temperature: goat anti-Rabbit IgG peroxidase (Sigma #A0545) and goat anti-mouse IgG peroxidase (Sigma #A0168). Each membrane was incubated with WesternBrightTM Quantum (advanstar, K-12042-D20) solution for two minutes, followed by acquisition of a chemiluminescence image using an iBright imaging system (ThermoFisher).

### In-cell interaction validation with LuTHy

LuTHy-BRET assays were performed as described previously (Trepte et al., 2018). In brief, cells were reverse transfected using linear polyethyleneimine (25 kDa, Polysciences 23966) with LuTHy constructs and cells were subsequently incubated for 48 h. Previously generated LuTHy control vectors expressing only NL (Addgene #113442) or PA-mCit (Addgene #113443) were used as background controls. Live cell BRET measurements were carried out in flat-bottom white 96-well plates (Greiner, 655983) with 24 PPIs per plate (each PPI in triplicate). Infinite® microplate readers M1000 or M1000Pro (Tecan) were used for the readout with the following settings: fluorescence of mCitrine recorded at Ex 500 nm/Em 530 nm, luminescence measured using blue (370–480 nm) and green (520–570 nm) band pass filters with 1,000 ms integration time. BRET ratios were calculated by dividing the background corrected luminescence intensity at 520-570 nm by the intensity obtained at 370-480 nm and subsequent donor bleed-through subtraction from NL only expressing wells.

## Acknowledgements

This research was supported by CHDI Foundation (RJH, CHA, EEW and CS), the Huntington’s Disease Society for America (ESR), and in part by the intramural research program of the NIH (SML, JCD). The Structural Genomics Consortium is a registered charity (no: 1097737) that receives funds from Bayer AG, Boehringer Ingelheim, Bristol Myers Squibb, Genentech, Genome Canada through Ontario Genomics Institute [OGI-196], EU/EFPIA/OICR/McGill/KTH/Diamond Innovative Medicines Initiative 2 Joint Undertaking [EUbOPEN grant 875510], Janssen, Merck KGaA (aka EMD in Canada and US), Pfizer and Takeda.

## Data Availability

Cryo-EM maps can be downloaded at EMDB-28767 and EMDB-28766. All constructs generated in this study have been deposited in Addgene and are detailed in Appendix 1.

## Contributions

RJH, CHA conceptualisation; MGA, RJH, JCD, PL, AH, AS, RC, ESR, CS, MA methodology; MGA, RJH, JCD, ESR, CS, MA analysis; MGA, RJH, JCD, ESR, CS, MA investigation; MGA, RJH, JCD, ESR, CS, MA manuscript preparation; MGA, RJH, JCD, ESR, CS, EW, SML, CHA manuscript review and editing.

## Abbreviations

HTT: huntingtin
HAP40: huntingtin associated protein 40 kDa
CTD: C-terminal domain
N-terminal domain
LuTHy: luciferase two-hybrid assay
cryo-EM: cryo-electron microscopy

