## Supplementary Information for "Expanding the Huntington’s disease research toolbox; validated subdomain protein constructs for biochemical and structural investigation of huntingtin"

*Corresponding Authors

### Abstract

Huntington's disease is characterised by CAG expansion in the huntingtin gene above a critical threshold of ~35 repeats, resulting in polyglutamine expansion of the huntingtin protein (HTT). The biological role of wildtype HTT and the associated mechanisms of disease pathology caused by expanded HTT remain incompletely understood, in part, due to challenges characterising interactions between HTT and putative binding partners. Here we describe a biochemical toolkit of rationally designed, high-quality recombinant HTT subdomains; one spanning the N-terminal HEAT and bridge domains (NTD) and the second spanning the C-terminal HEAT domain (CTD). Using biophysical methods and cryo-electron microscopy, we show these smaller subdomains are natively folded and can associate to reconstitute a functional full-length HTT structure capable of forming a near native-like complex with 40 kDa HTT-associated protein (HAP40). We report biotin-tagged variants of these subdomains, as well as full-length HTT, that permit immobilisation of each protein for quantitative biophysical assays without impacting protein quality. We demonstrate the CTD alone can form a stable complex when co-expressed with HAP40, which can be structurally resolved. The CTD-HAP40 complex binds the NTD, with a dissociation constant of approximately 10 nM as measured by bio-layer interferometry. We validate the interaction between the CTD and HAP40 using a luciferase two-hybrid assay and use subdomain constructs to demonstrate their respective stabilization of HAP40 in cells. These open-source biochemical tools will enable the wider HD community to study fundamental HTT biology, discover new macromolecular or small-molecule binding partners and map interaction sites across this very large protein.

### Keywords

Huntington disease, neurodegeneration, protein purification, protein structure, cryo-electron microscopy, protein-protein interaction.

*To whom correspondence should be addressed:

Rachel J. Harding


Cheryl H. Arrowsmith


**Table S1: Cryo-EM data collection, refinement and validation statistics.**

| | Full-length HTT-HAP40 (EMDB-28767) | CTD-HAP40 (EMDB-28766) |
|---|---|---|
| Data collection and processing | | |
| Magnification | 100,000 | 100,000 |
| Voltage (kV) | 200 | 200 |
| Electron exposure (e–/Å$^2$) | 51.3 | 51.3 |
| Defocus range (µm) | -2.0 to -0.4 | -2.0 to -0.4 |
| Pixel size (Å) | 0.81 | 0.81 |
| Symmetry imposed | C1 | C1 |
| Initial particle images (no.) | 112,717 | 986,229 |
| Final particle images (no.) | 64,597 | 134,849 |
| Map resolution (Å) | 3.3 | 3.2 |
| FSC threshold | 0.143 | 0.143 |

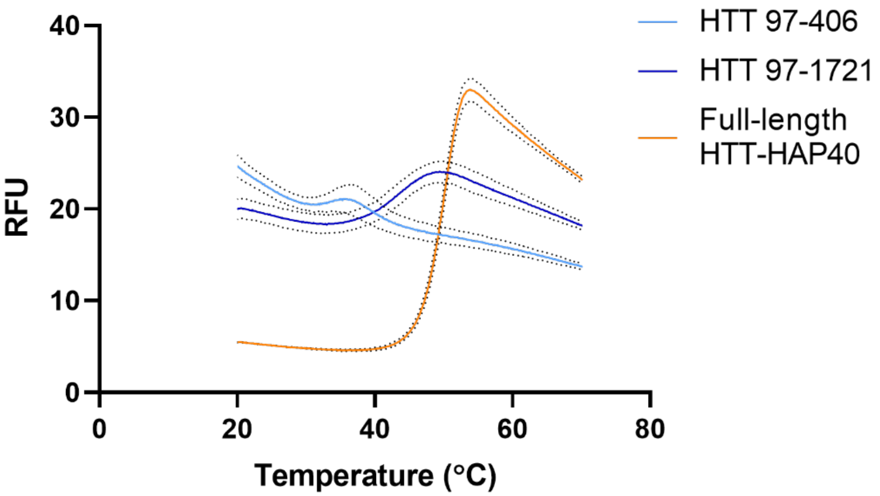


**Figure S1: N-terminal HTT fragments comprising amino acids 97-406 and 97-1721 are not viable protein constructs as determined by DSF.** Samples were run at a final concentration of 0.1 mg/mL protein and 5X Sypro Orange in buffer containing 20 mM HEPES pH 7.4, 300 mM NaCl, 2.5% (v/v) glycerol, 1 mM TCEP. Temperature ramp rate of 0.02°C / sec was used. Dotted lines indicate 1 S.D. from mean triplicate values.

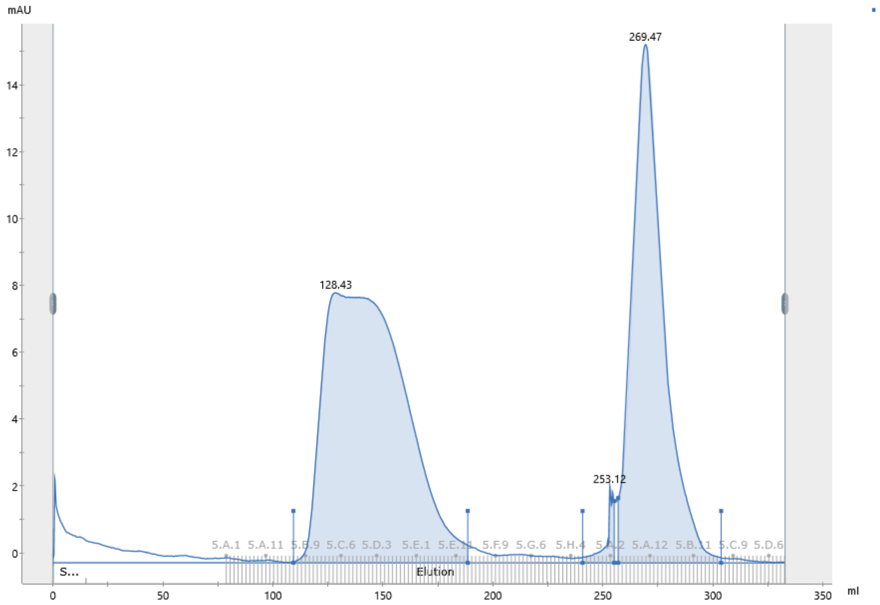


**Figure S2: Size exclusion chromatography profile of HTT aa. 97-1721 N-terminal HTT fragment indicates the sample is aggregated.** The protein elutes at the void volume of the column (~120 mL) and a monodisperse peak was not resolved. A HiLoad 26/600 Superdex 200 column (Cytiva) was used at a flow rate of 2 mL/min.

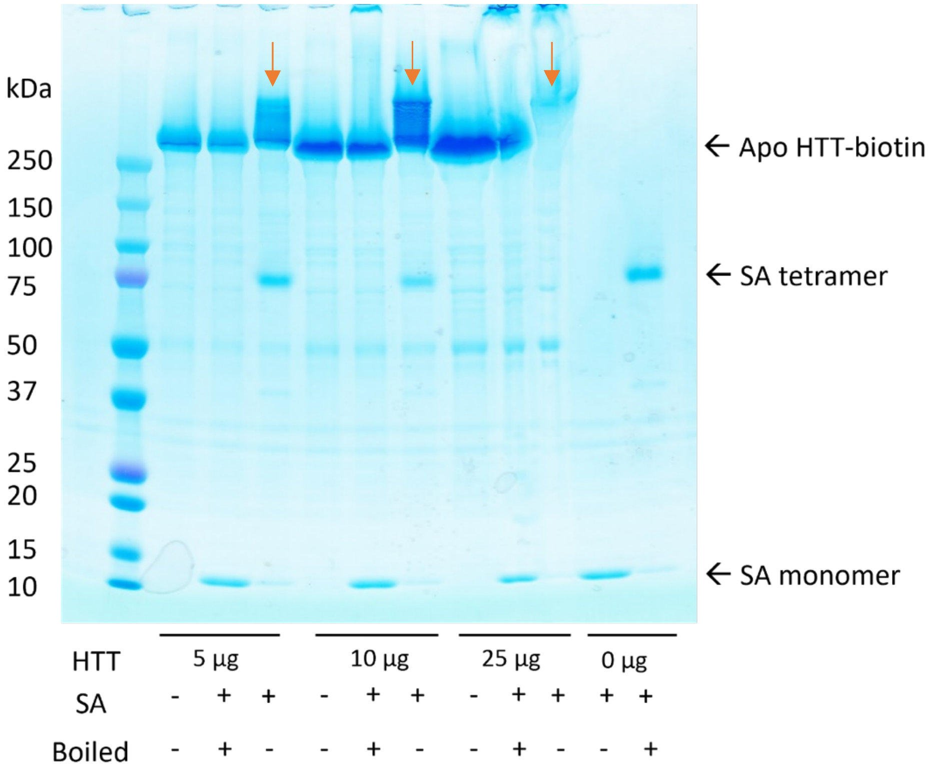


**Figure S3: Representative streptavidin gel shift assay to confirm the presence of biotin tag on Avi tagged HTT constructs.** Various concentrations of biotinylated HTT were incubated with 2 µg streptavidin (SA). SA retains its folded state as a tetramer and its ability to bind biotin when loaded onto an SDS-PAGE without boiling the sample. Increasing the amount of biotinylated HTT incubated with SA results in reduced band intensity corresponding to the SA tetramer, and high molecular weight HTT-SA complex bands (orange arrows), indicating that HTT is biotinylated and binds SA.

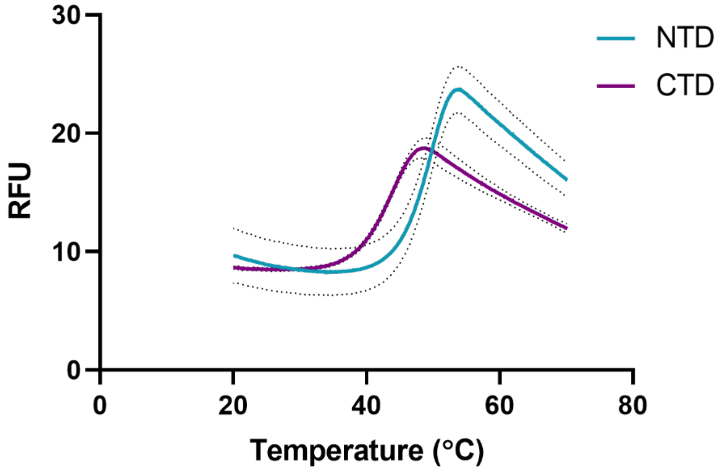


**Figure S4: Unprocessed differential scanning fluorimetry (DSF) profiles of purified HTT N-terminal HEAT and bridge domain (NTD) and C-terminal HEAT domain (CTD).** Both proteins display good dynamic range of fluorescent signal and sharp transitions, indicating a change from folded to unfolded states upon heating. Samples were run at a final concentration of 0.1 mg/mL protein and 5X Sypro Orange in buffer containing 20 mM HEPES pH 7.4, 300 mM NaCl, 2.5% (v/v) glycerol, 1 mM TCEP. Temperature ramp rate of 0.02°C / sec was used. Dotted lines indicate 1 S.D. from mean triplicate values.

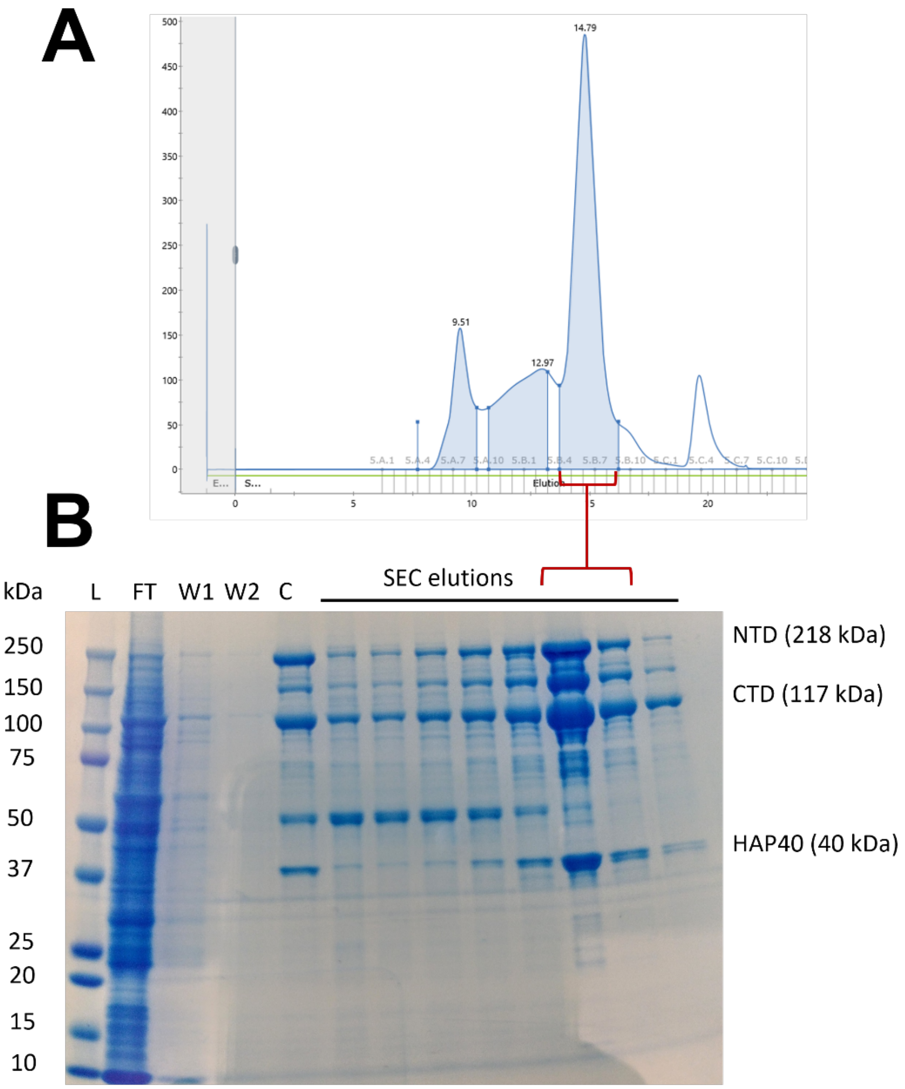


**Figure S5: Purification of co-expressed NTD, CTD and HAP40.** *A*, Size-exclusion chromatography profile of crude protein isolated from 4 L Sf9 culture through FLAG affinity purification. 1 mL crude concentrated protein was loaded onto a Superdex 200 10/300 column (Cytiva) and eluted at a flow rate of 0.5 mL/min. *B*, SDS-PAGE of protein fractions from FLAG affinity purification and SEC. L, ladder; FT, FLAG purification flow-through; W1, first wash; W2, second wash; C, crude eluate.

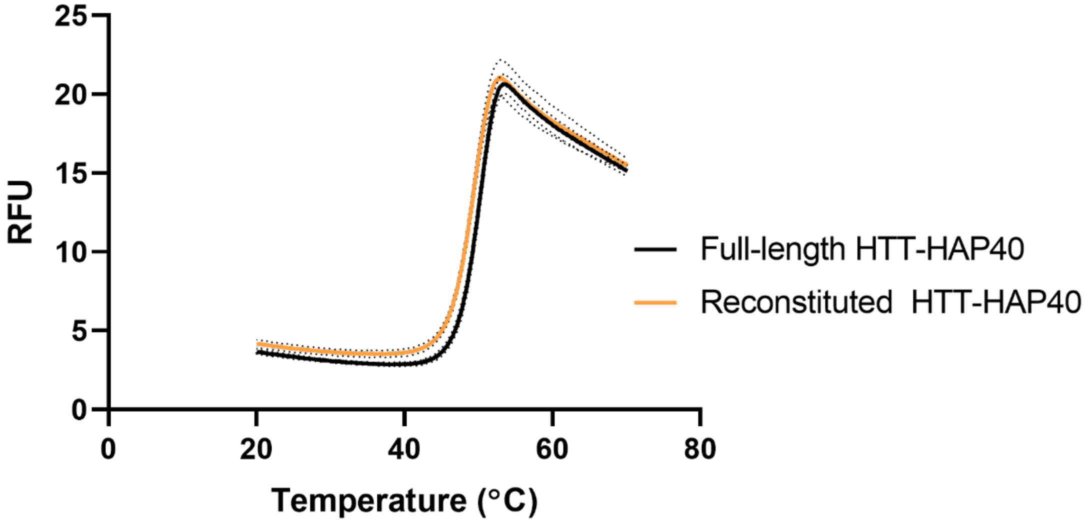


**Figure S6: Unprocessed differential scanning fluorimetry (DSF) profiles of full-length HTT-HAP40 vs reconstituted HTT through generated through co-expression of NTD, CTD and HAP40 domains.** Samples were run at a final concentration of 0.1 mg/mL protein and 5X Sypro Orange in buffer containing 20 mM HEPES pH 7.4, 300 mM NaCl, 2.5% (v/v) glycerol, 1 mM TCEP. Temperature ramp rate of 0.02°C / sec was used. Dotted lines indicate 1 S.D. from mean triplicate values.

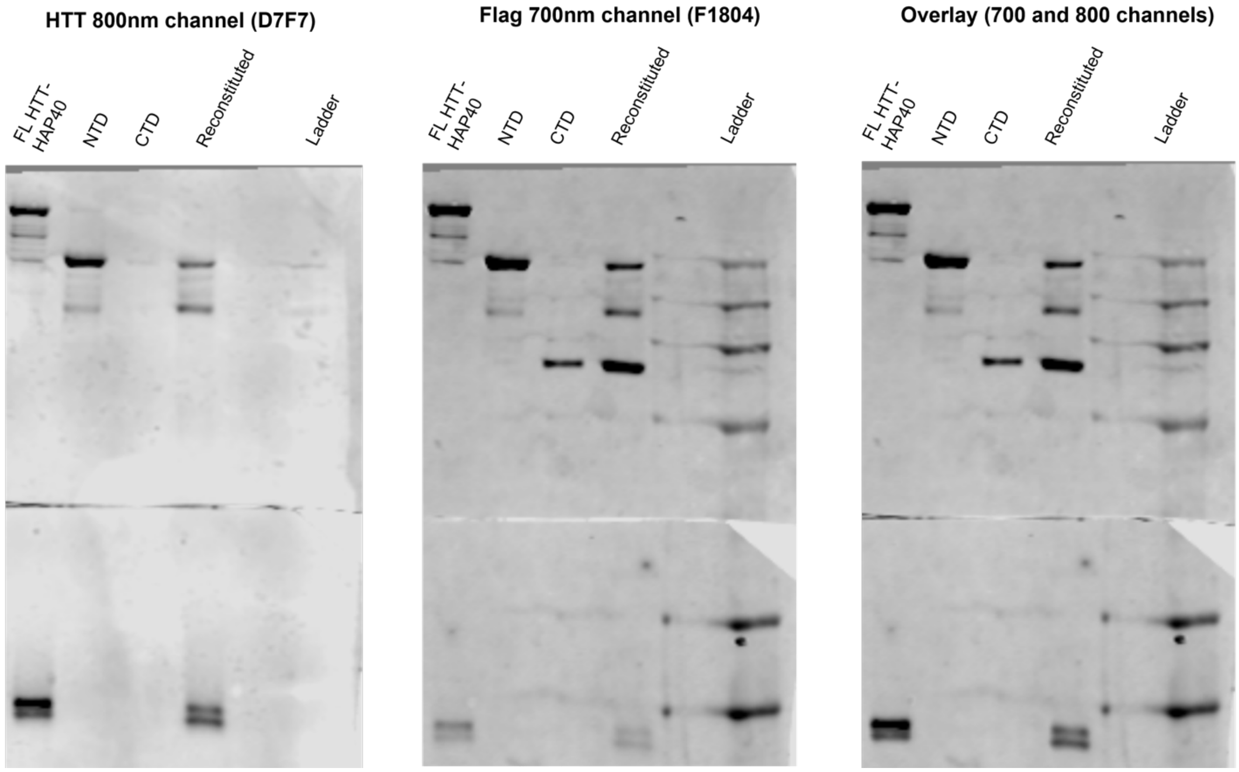

**Figure S7: Complete western blots of full-length HTT-HAP40, NTD and CTD subdomains, and reconstituted HTT-HAP40 co-expression using the NTD and CTD.**

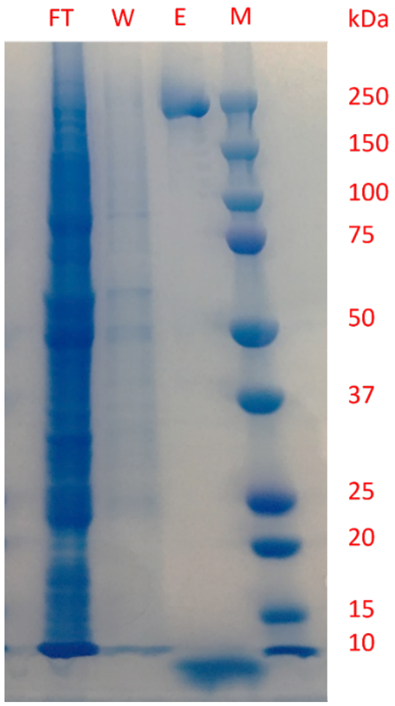


**Figure S8: Expression of HAP40 is not viable when co-expressed with NTD as determined by SDS-PAGE of FLAG affinity purification of the coexpressed constructs.** *FT*, flow-through; *W*, wash; *E*, elution; *M*, marker.

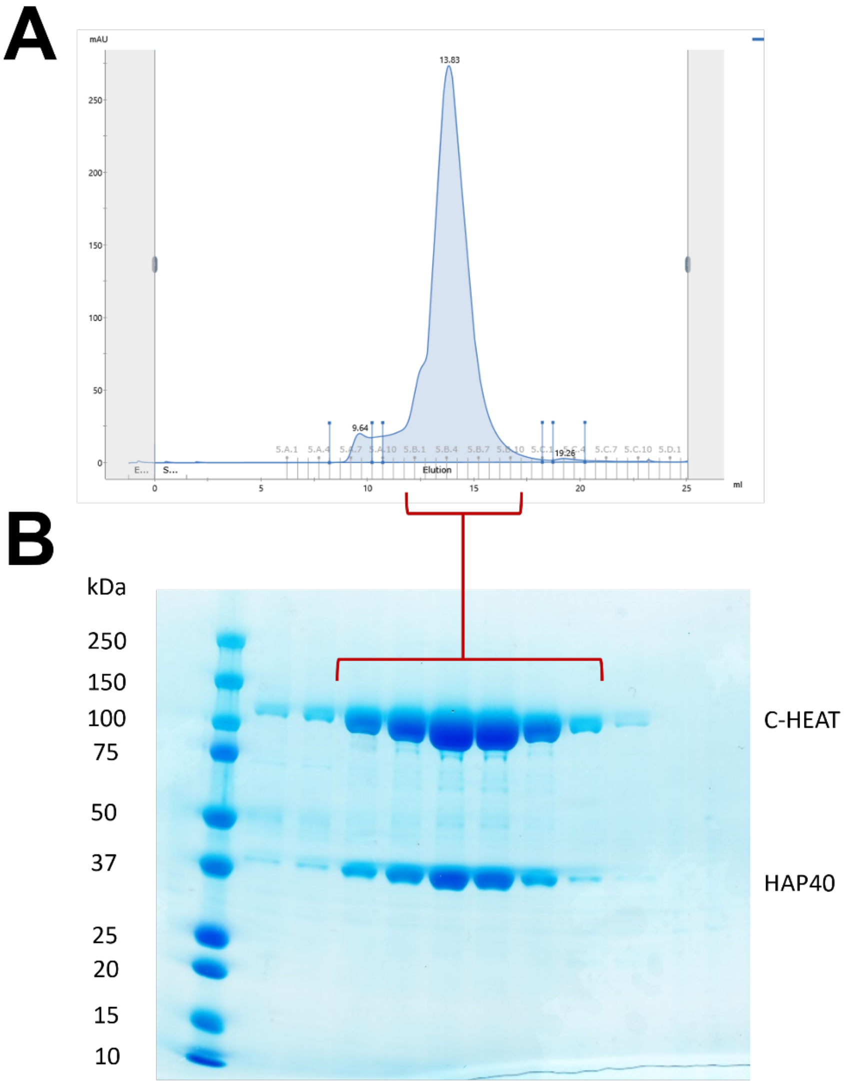


**Figure S9: Expression of HAP40 is viable when coexpressed with the CTD.** *A*, Size-exclusion chromatography profile of crude protein isolated from 4 L Sf9 culture through FLAG affinity purification. 1 mL crude concentrated protein was loaded onto a Superdex 200 10/300 column (Cytiva) and eluted at a flow rate of 0.5 mL/min. *B*, SDS-PAGE analysis of fractions collected from size-exclusion chromatography. Indicated fractions from monodisperse protein peak were pooled, concentrated and flash frozen.

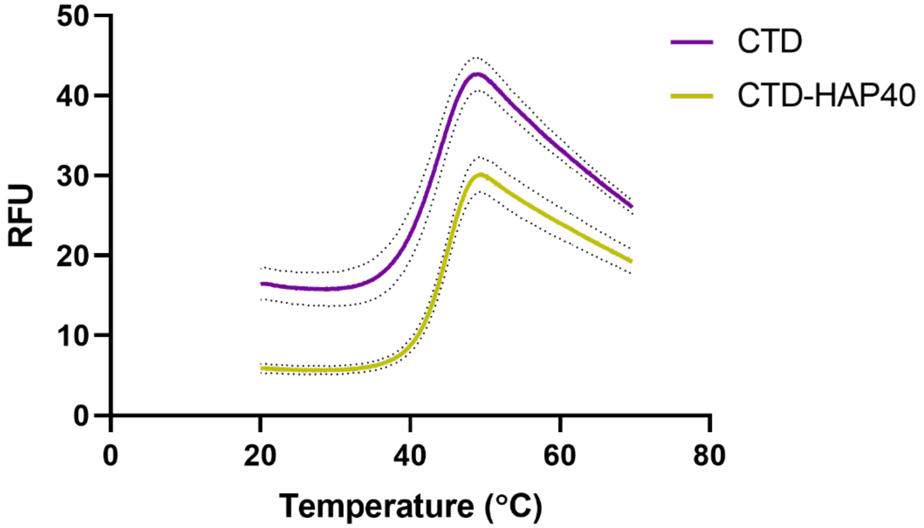


**Figure S10: Unprocessed differential scanning fluorimetry (DSF) profiles of apo-CTD compared with co-expressed CTD-HAP40.** Samples were run at a final concentration of 0.1 mg/mL protein and 5X Sypro Orange in buffer containing 20 mM HEPES pH 7.4, 300 mM NaCl, 2.5% (v/v) glycerol, 1 mM TCEP. Temperature ramp rate of 0.02°C / sec was used. Dotted lines indicate 1 S.D. from mean triplicate values.

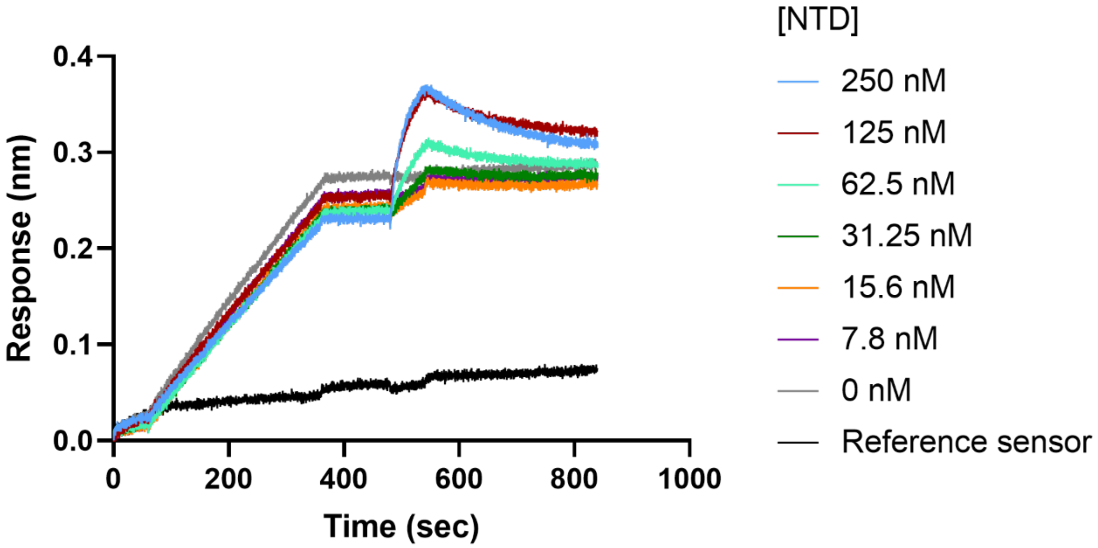


**Figure S11: Complete unprocessed BLI sensorgrams measuring the kinetics of NTD binding to immobilised biotinylated CTD.** Streptavidin biosensors (ForteBio) were loaded with 1 µg/mL biotinylated CTD ($t_{60} – t_{360}$), followed by a 120 second baseline equilibration ($t_{360} – t_{480}$). Various concentrations of NTD were incubated with the sensors during a 60 s association phase ($t_{480} – t_{540}$) followed by a 300 s dissociation phase in buffer ($t_{540}$ – $t_{840}$). Reference sensors without immobilised CTD were incubated with 250 nM NTD. Signal from reference and 0 nM NTD sensors were subtracted from experimental wells.
